# Genomic sequencing to detect cross-breeding quality in dogs: an example studying disorders in sexual development

**DOI:** 10.1101/2024.08.07.606952

**Authors:** Luciana de Gennaro, Matteo Burgio, Giovanni Michele Lacalandra, Francesco Petronella, Alberto L’Abbate, Francesco Ravasini, Beniamino Trombetta, Annalisa Rizzo, Mario Ventura, Vincenzo Cicirelli

**Author notes:** These authors equally contributed to this work (Luciana de Gennaro, Matteo Burgio). Corresponding authors (M.V.); (V.C.).

## Abstract

**Background:** Disorders of Sexual Development (DSD) in dogs, similar to humans, arise from irregularities in genetic determinants, gonadal differentiation, or phenotypic sex development. The French Bulldog, a breed that has seen a surge in popularity and demand, has also shown a marked increase in DSD incidence. This study aims to characterize the genetic underpinnings of DSD in a French Bulldog named Brutus, exhibiting ambiguous genitalia and internal sexual anatomy, and to explore the impact of breeding practices on genetic diversity within the breed.

**Methods:** We utilized a comprehensive approach combining conventional cytogenetics, molecular techniques, and deep sequencing to investigate the genetic profile of Brutus. The sequence data were compared to three other male French Bulldogs genome sequences with typical reproductive anatomy, including Brutus’s father, and the canine reference genome (CanFam6).

**Findings:** Our findings revealed a 22% mosaicism (78, XX/77, XX), the absence of the SRY gene, and the presence of 43 unique Single Nucleotide Variants (SNVs) not inherited from the father. Notably, the Run of Homozygosity (ROH) analysis showed Brutus has a significantly higher number of homozygous segments compared to other Bulldogs, with a total length of these fragments 50% greater than the average, strongly suggesting this dog is the product of the mating between siblings. While no direct causative genes for the DSD phenotype were identified four candidate loci warranting further investigation were highlighted.

**Conclusions:** Our study highlighted the need for a better annotated and curated reference dog genome to define genes causative of any specific phenotype, suggests a potential genetic basis for the DSD phenotype in dogs, and underscores the consequences of uncontrolled breeding practices in French Bulldogs. These findings highlight the importance of implementing strategic genetic management to preserve genetic health and diversity in canine populations.

## Introduction

Physiological sexual differentiation in humans as in dogs, depends on genetic determinants, gonadal differentiation, and development of the phenotypic sex. An irregularity in any of these three steps can lead to Disorders in Sexual Development (DSD) (1,2). Dog has been used as an experimental animal model for DSD to study human sexual development disorders, because of its high similarity with humans (3,4). These conditions are considered rare in the canine species and their variability in phenotypic expression with ambiguous external genitalia makes the diagnosis and a unique classification of intersex conditions difficult (3). Following the human classification (5) there are three major categories: (1) DSD caused by an abnormal set or structure of sex chromosomes, called sex chromosome DSD; (2) DSD with a normal female set of sex chromosomes, called XX DSD and (3) DSD with a normal complement of male chromosomes, XY DSD (2,4). The first group includes all those subjects whose karyotype shows numerical variations of chromosomes such as monosomies of the X chromosome, trisomies XXX or XXY, and chimerism phenomena in which subjects show cells with both XX and XY sex chromosomes at the same time.

Despite numerous studies, the chromosomal background of these anomalies in dogs remains unclear (6) mostly because karyotyping in dogs presents a significant challenge due to the high chromosome count (2n = 78) and the acrocentric nature of all predominantly small autosomes (7). Only the sex chromosomes are metacentric and easily distinguishable from the autosomes. Easily identifiable mutations, such as those involving sex chromosome aneuploidies primarily associated with DSDs are X monosomy (8), X trisomy (9), and XXY sex chromosomes (10).

Despite extensive analyses, no causative point mutations have been found in specific candidate genes to explain these disorders. However, a few studies have shown a link between XX DSD and mutations such as copy number variations, duplications, and substitutions in SOX9 (Sry-box containing gene 9 or Sry-like HMG box) gene, a key gene having a role in testis induction in vertebrates (11). The SRY gene, essential for male development, provides further insights into DSD types by working with the SOX9 gene to produce testosterone and Anti-Müllerian hormone (AMH), leading to male genitalia formation (2,12). Recent research has suggested an association between PAD16 gene variants and XX DSD in certain dog breeds, highlighting the complex and breed-specific nature of genetic variations in DSD phenotypes (13). DSDs have been detected in more than 40 dog breeds with an increased incidence in American Cocker Spaniels, English Cocker Spaniels, Kerry Blue Terriers (14), American Staffordshire terriers and French Bulldogs (2). XX DSDs are the most frequently seen disorders with an increase in incidence in recent years in the French Bulldog. This is probably due to uncontrolled mating that responds to the largely increased demand of the market. French Bulldog represents the most popular breed in the UK in 2018 (2,15,16). Recent data on DSDs in dogs have highlighted the significant increase over the years of this disorder in the French Bulldog breed (2). This increase in the number of cases clearly outlines a stable growth trend that suggests the need to investigate the underlying causes of this process for a clearer view of the pathology.

For this reason, we used a multipronged approach to deeply characterize an 11-month-old French bulldog, named Brutus, with ambiguous external genitalia consisting of a vulva containing a penis-like structure and internal genitalia displaying two testicles connected to a complete uterus. Using conventional and molecular cytogenetics, deep sequencing, and molecular biology techniques, we deeply investigated the genetic profile of the phenotypic condition associated with our sample. Sequence data were compared to three other male French bulldogs with regular reproductive activities, including Brutus’ father, and canine reference genome (CanFam6). We found the presence of 22% mosaicism 78, XX/77, XX, the absence of the SRY gene, and 43 unique not inherited from the father SNVs compared to the other dogs. Of note, eight of these SNVs are clustered within an 800 bp region on the X chromosome, upstream of the spermiR sequence, which is critical for spermatogenesis in some mammals, suggesting that these SNVs may play a regulatory role in male reproductive development also in dogs. This finding highlights the necessity of further studies on the function of this region in dogs.

Furthermore, the run of homozygosity (ROH) analysis revealed that Brutus has the highest number of homozygous segments than the others, leading to a total length of these homozygous fragments 50% higher than the average of other individuals from different breeders. Although this work did not allow us to find a direct correlation with the DSD phenotype, we were able to highlight RSPO1, SOX9, and SRD5A2 as candidate genes that need further study. Noteworthy, the sequencing approach allowed us to highlight how the uncontrolled cross-breeding carried out by Bulldog breeders is currently having genotypic and consequently phenotypic effects. This underscores the need for such an approach to properly plan genetic management in canine populations.

## Materials and Methods

### FISH Experiments

Canine lymphocytes derived from Brutus’s peripheral blood sample were stimulated with phytohemagglutinin (PHA) and used to prepare the metaphase spreads with the standard procedures (17).

Cytogenetic analyses were performed by fixing metaphase spreads onto slides and then incubated at 90°C for 1h30′ to contribute to sample fixation and dehydration. 0.005% pepsin/HCl 0.01 M treatments were performed to eliminate cytoplasm proteins for better hybridization rates.

Subsequent treatments in PBS 1X, MgCl2 0.5M, 8% paraformaldehyde, and 70%/90%/absolute alcohols allowed a proper stabilization, fixation, and dehydration of metaphases and nuclei DNA molecules.

In situ fluorescence hybridization (FISH) experiments were performed on nuclei and metaphases using probes from a female canine BAC CHORI-82 library (https://bacpacresources.org/library.php?id=253) and human RP11 BAC library (Table 1).

**Table 1.** Probes used for the FISH experiments. The mapping for the BAC probes derived from the canine CH82 library is based on the CanFam6 reference genome, while the BAC probe from the human RP11 library is mapped to the hg38 reference genome.

| PROBE | MAPPING |
| --- | --- |
| CH82-201N14 | chrX:92883400-93054893 |
| CH82-509B23 | chr5:30780182-30905463 |
| CH82-253P13 | chr23:26062226-26251145 |
| CH82-26I8 | chr9:9505916-9667419 |
| RP11-400O10 | chrY:2724275-2921156 |

DNA extraction from selected BACs has been done with the Biorad Quantum Prep Plasmid Miniprep Kit. FISH experiments were performed essentially as previously described (18): two hundred nanograms of the DNA probe, labeled by nick-translation with Cy3-dUTP or fluorescein-dUTP, were precipitated by ion-exchange alcohol precipitation with canine Cot DNA and finally denatured for 2 minutes at 70°C and hybridized at 37°C overnight. Post-hybridization washing was at 60°C in 0.1× SSC (three times, high stringency). The hybridization with the human probe was instead performed at 72°C for 2 minutes, and post-hybridization washes were carried out under low stringency conditions, specifically three washes at 38°C with 2x SSC. The slide was then stained with DAPI, producing a Q-banding pattern. The fluorescence signals coming from Cy3, Fluorescein, and DAPI were detected separately with specific filters using a Leica DMRXA epifluorescence microscope equipped with a cooled CCD camera (Princeton Instruments) and recorded as grayscale images. Finally, Adobe Photoshop™ software was used for image pseudo-colorization and merging of the acquired images.

### Genomic extraction

Genomic DNA of all the samples was extracted starting from blood samples with the DNA Blood mini-kit (Qiagen Inc.), following the manufacturer’s protocol. The Nanodrop spectrophotometer measured the DNA concentrations.

### PCR

To investigate the presence of SRY in Brutus a PCR using primers specific for canine SRY (Forward: 5’ AAGGCCACGGCACAGAAAAGTCAC and Reverse: 5’ AAGAAGCGTCAGCGGACATCTGTG from (19) was performed using iProof HF Master Mix (BioRAD). Amplification was carried out in 25 μL reactions with 1 × PCR MasterMix, 0.5 μm forward and reverse primers, and 50 ng of genomic DNA. The reaction was then cycled with the following conditions: initial denaturation at 98°C for 3 min, then 30 cycles at 98°C for 10 sec, 70°C for 30 sec, and 72°C for 20 sec; final extension was at 72°C for 7 min.

### Sequencing analysis: Variant calling and ROH estimation

The extracted DNA was sent to the CD Genomics Institute for DNA QC, Library Preparation, and NovaSeq6000 S4 Sequencing (PE152, 30X).

Raw reads were aligned to the last dog reference genome available (Dog10K_boxer_Tasha/canFam6, Oct. 2020) using BWA-MEM algorithm (v.0.7.17) with the default parameters (https://bio-bwa.sourceforge.net), and we removed PCR duplicates using the Picard MarkDuplicates tool (version 3.1.1., http://broadinstitute.github.io/picard/). Variant calling analysis was performed using GATK (v4.3.0.0) (20) with “*HaplotypeCaller*“ and “*-ERC* GVCF“. After combining (“CombineGVCFs”) the gVCFs of the individual samples, the merged file was then genotyped (“GenotypeGVCFs”) and filtered only for SNP-type variants (“*SelectVariants*” and “*--select-type-to-include* SNP” options). Using “*VariantFiltration*“ of GATK with the “*--filter-expression*” option, we discarded SNVs with a “DP“ value (read depth) lower than the 75th percentile of the DP values, and the SNPs with QD < 2.0, FS > 60.0, MQ < 56.0, SOR > 3.0, MQRankSum < -12.5, ReadPosRankSum < -8.0.

By evaluating only the positions in which there was a call in all the samples, only the SNVs in which a specific variant is present only in Brutus and never, even in heterozygosity, in the other samples.

We downloaded from UCSC Genome Browser the complete SNPs database (dbSNPs), based on ‘canFam4’ genome release (https://genome.ucsc.edu/cgi-bin/hgTables?hgsid=2314426054_ivCEAdUZP0HdC5ZSWlzUb5F20Rzu) and we converted all SNPs position to the ‘canFam6’ release, the one used for generating the BAM files. We used the converted dbSNPs to filter out common SNPs in our samples. Finally, SnpEff (v5.2) (https://pcingola.github.io/SnpEff/) software was used for annotating the retained list of variants.

To quantify genomic inbreeding, we measure the proportion of an individual’s DNA inherited from recent common ancestors, providing an estimate of the percentage of the individual’s genome that is inbred. Runs of Homozygosity (ROH) were identified using the *--homozyg* function in PLINK v1.9 considering only genomic segments extending more than 1000 kb (*--homozyg-kb*) and with at least 100 SNPs (*--homozyg-snp*). The resulting ROH segments were analyzed to determine the genomic inbreeding coefficient (F_ROH), which represents the proportion of the autosomal genome (total length of autosomal chromosomes = 2,212,284 kb) covered by these segments.

After analyzing the ROHs, we identified the genes in Brutus’ homozygous segments. The analysis was performed in R using the *foverlaps* function from the *data.table* library, following a liftover to canFam4 to utilize the most comprehensive RefSeq track available. We used canFam4 instead of the last release canFam6, because of the highest accuracy in gene annotation of the former. The obtained gene list was analyzed using the ToppGene portal (https://toppgene.cchmc.org/), as described in (21), a comprehensive platform for gene list enrichment analysis and candidate gene prioritization based on functional annotations and protein interaction networks. The ToppFun function was specifically utilized to detect functional enrichment of genes based on various ontologies (GO, pathway), mouse phenotype, literature cocitation, and other features.

## Results

Brutus, at the clinical examination, showed a generally good state of nutrition. The external genitalia showed a normal vulva in terms of position, shape, and size, the vulvar lips showed inflammation and excoriation of the skin part caused by the patient’s continuous licking of the area. The labia could not close the vulvar opening due to the presence and protrusion of the penile structure. The penis, containing the penile bone, showed a dry, inflamed, and eroded surface due to the lack of skin covering and protection. There were no abnormalities in the course of the internal urethra and the external part of the organ connected to the external sphincter located on the dorsal face of the body of the penis. The internal genitalia, investigated by second-level diagnostic means (ultrasound and computed tomography) showed the presence of two compact intra-abdominal parenchymatous structures connected to two cavitary organs referable to uterine horns (S1 Text). Following laparoscopic surgery to remove the intra-abdominal organs, histological analysis showed testicular parenchyma presenting seminiferous tubules with undeveloped germinal epithelium and Sertoli cells with nucleated nuclei and no mature spermatozoa (Fig 1).

**Fig 1.**
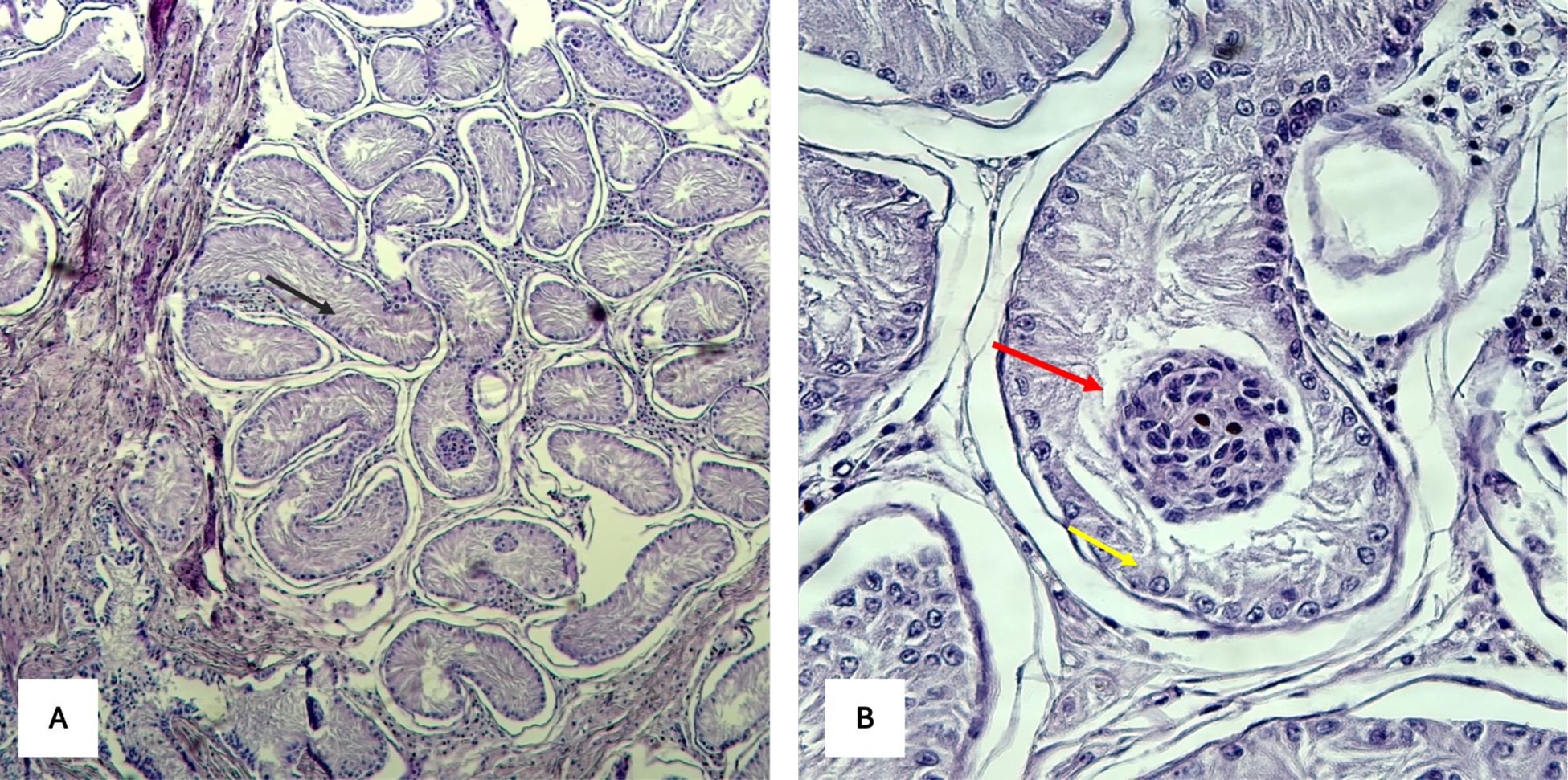
Histological analysis of the testicular parenchyma removed during laparoscopic surgery. The images display tissue sections stained with hematoxylin and eosin (H&E), revealing the structural and cellular characteristics of the testicular tissue. (A) Seminiferous tubule with undeveloped germinal epithelium (black arrow); (B) Sertoli cells with nucleated nuclei (yellow arrow), immature germ cell aggregates in the lumen (red arrow).

Additionally, seven slides with metaphase preparations were obtained from the blood of Brutus, stained with DAPI, and visualized under a fluorescence microscope to evaluate the chromosomal structure and perform a classical karyotype analysis. All observed metaphases, obtained from the lymphocyte culture, were karyotyped using standard karyotype as a reference (7) and showed the presence of two metacentric chromosomes X and the absence of the Y chromosome. To confirm the identity of these large submetacentric chromosomes present in all metaphases, we performed a FISH experiment using a dog-specific probe (CH82-201N14, chrX:92883400-93054893 CamFam6 reference genome). This probe showed different signal intensities on interphase nuclei and the two homologous chromosomes (Fig 2A), showing a heterozygous state for this genomic copy number variant.

**Fig 2.**
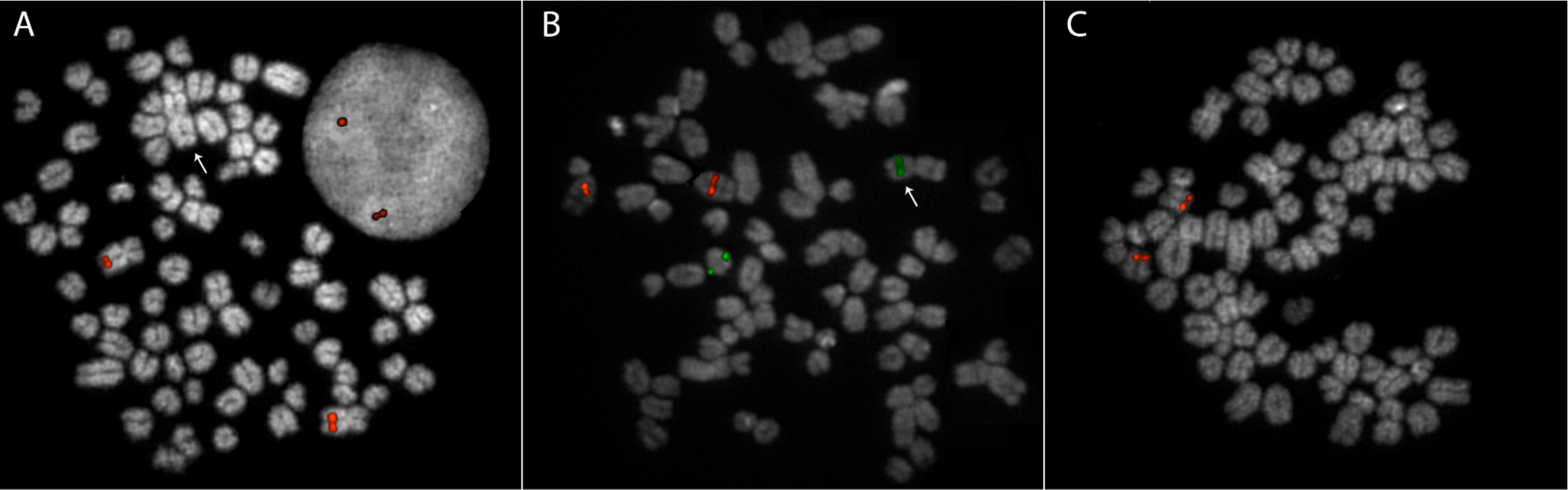
FISH analysis on Brutus using different probes. (A) FISH with an X-specific probe (CH82-201N14), showing both metaphase chromosomes and interphase nuclei to highlight the polymorphism of this region. (B) FISH using probes for chromosome 5 (CH82-509B23; red) and chromosome 23 (CH82-253P13; green). (C) FISH with a SOX9-specific probe (CH82-26I8).

The chromosome count was performed and in 18 out of 82 metaphase spreads, we observed 77 chromosomes with an additional non-acrocentric chromosome lacking the chromosome X-specific signal (Fig 2A). A FISH experiment using two specific probes for chromosomes 5 and 23, CH82-509B23, and CH82-253P13 respectively, was performed to check whether the additional metacentric chromosome found in some metaphases was the result of a Robertsonian translocation like the one previously reported in another case of Sex Reversal Dog (22). Of note we detected the presence of a chromosome 23 probe on the additional metacentric, showing a different Robertsonian fusion than the previously reported (Fig 2B).

The conventional and molecular cytogenetic analysis was also performed on the other 3 healthy control dogs analyzed, including Brutus’ father, and in all cases a normal 78, XY karyotype was found (S1 Fig).

To investigate the presence of the SRY gene, an additional FISH experiment with a human-specific probe (RP11-400O10) for the SRY gene was used on the 4 dogs we were testing. None of the experiments was successful, likely due to the high divergence existing between the target (dog genome) and probe (human source). We further investigate the presence of the SRY gene in Brutus by performing a PCR using primers canine-SRY-specific (19). 271 bp expected amplification product was obtained only in Brutus’ father and in the other two controls (Tauro and Bufalo), confirming what was already karyotypically hypothesized that our case with DSD is also SRY negative (S2 Fig).

The SOX9 gene which is often associated with DSD in humans (23) was also investigated using the CH82-26I8 probe specific for this region present on chr 9 (24), however, no significant deletion or duplication was highlighted (Fig 2C).

The alignment and variant calling of the sequenced genomes were performed using CanFam6 as the reference genome. The decision to use the genome of the Boxer breed as a reference is supported by phylogenetic (25) and cluster (26) analyses in the literature, which demonstrated the highest correlation between this genome and the French Bulldog. This approach ensures that the reference genome is closely related to the genetic background of our samples, thereby increasing the accuracy and relevance of the alignment and variant calling results. Following the annotation performed on the variant calling output (see Materials and Methods), we identified a total of 43 SNVs that are unique to the Brutus genome. All these variants identified have been classified, after SNPeff annotation, as having a putative impact level of “modifier” (S1 Table). 20 out of the 43 identified SNVs are on autosomal chromosomes (2, 8, 14, 17, 19, and 20), 9 on the X chromosome, and 14 on mitochondrial DNA. Notably, SNPs on chromosomes 19 and X in Brutus are both homozygous and identical to the boxer reference genome. These variants could be significant for DSD phenotype since they were never detected in the other healthy Bulldogs sequenced. The alleles detected in the other three bulldogs could then represent specific breed variations.

To assess the homozygosity patterns within our samples, we conducted a Runs of Homozygosity (ROH) analysis, focusing on genomic segments exceeding 1,000 kb and containing at least 100 SNPs. Our analysis revealed that the healthy dogs (Tauro, Bufalo, and Brutus’s father) exhibited an average of 159 ROH segments, with a mean total length of 303,390 kb. In contrast, Brutus significantly deviates from these average values; the ROH analysis for Brutus identified 298 homozygous segments, with a total length (681,002 kb) exceeding twice the average length.

The details of the localization, number of SNPs, and percentage of homozygosity for each ROH fragment can be found in S2 Table.

Considering the total length of autosomal chromosomes, which is 2,212,284 kb, the values of F_ROH (fraction of ROH) for Brutus are markedly higher than those observed in the other samples and higher than the expected value between the two siblings (0.25) (Table 2).

**Table 2.** ROH analysis results for the healthy control dogs (Tauro, Bufalo, and Brutus’s father) and Brutus. Each column presents the Identifier for each dog (SAMPLE), the total count of homozygous segments identified (NSEG); the cumulative length of all ROH segments per sample (KB), the average length, in kb, of ROH segments (KBAVG) and the average inbreeding coefficient (F_ROH).

| SAMPLE | NSEG | KB | KBAVG | F_ROH |
| --- | --- | --- | --- | --- |
| Brutus | 298 | 681002 | 2285.24 | 0.308 |
| Bufalo | 153 | 299507 | 1957.56 | 0.135 |
| FatherB | 170 | 318511 | 1873.6 | 0.144 |
| Tauro | 154 | 292152 | 1897.09 | 0.132<sub>274</sub> |

Within the 298 Brutus’s ROH segments, 593 genes were identified (according to the UCSC RefseqCurated database for CanFam4; S3 Table). Of the 593 genes found, only 508 were recognized and analyzed by the ToppGene portal. The excluded genes (“not found” in column 2 of S3 Table) were predominantly miRNAs and olfactory receptor genes. The results obtained from ToppGene were categorized into “Biological Process” (S4 Table), “Mouse Phenotype” (S5 Table), “Disease” (S6 Table), and “Pubmed” (S7 Table). Within each category, we searched for the keywords “Development”, “Sex” and “hermaphroditism/hermaphroditic”, to search for the most significant results associated with our phenotype of interest. We found 14 in Pubmed, 12 in Biological Process, 3 in Mouse Phenotype, and 1 in Disease (items highlighted in red in S4-7 Tables), involving a total of 81 genes (S8 Table). Among these, SRD5A2, RSPO1, and SOX9 genes merit further investigation in dogs, as they have already been associated with our phenotype of interest in other species according to Pubmed publications.

## Discussion

Using a multipronged molecular approach, we performed a comprehensive genetic analysis of a French Bulldog, named Brutus, with DSD and found significant information on this case’s chromosomal and genetic characteristics. Karyotypic analysis and molecular cytogenetics revealed the presence of two metacentric X chromosomes and the absence of the Y chromosome. The use of a specific probe targeting the X chromosome to confirm the sex chromosome configuration in Brutus revealed a notable difference in signal intensity compared to the control dogs. This variation indicates a heterozygous state for a genomic copy number variant on the X chromosome. The probe used is known to cover the region corresponding to the human RBMYA1 gene, which has been previously associated with male infertility in humans (24). The differential signal intensity observed in Brutus suggests that this genomic variant could be influencing the expression of genes linked to fertility. We confirmed the absence of the Sex Determining Region Y (SRY) in Brutus by PCR using canine-specific SRY primers. This chromosomal anomaly is consistent with findings from similar cases of sex reversal in dogs (15).

Additionally, we detected in 82 metaphases analyzed, approximately 22% 78, XX/77, XX mosaicism. Using both classical and reiterative FISH experiments, we identified an additional metacentric in the metaphase spread with 77 chromosomes as a result of a Robertsonian fusion between the 23 and an unidentified chromosome. Unlike a previously reported DSD case in which the translocation involved chromosomes 5 and 23 (22), our results did not show the involvement of chromosome 5, suggesting a different chromosomal partner in our translocation event. This highlights the genetic complexity of the DSD phenotype and supports the possibility that the development of DSD, although not related to a specific Robertsonian translocation, could somehow be associated with rearrangements of chromosome 23 that could potentially affect the regulatory sequences of genes located on chromosome 23 (FOXL2 and CTNNB1) involved in reproductive development (22).

The absence of the SRY gene in Brutus, confirmed by PCR, identified this dog as an SRY-negative DSD phenotype. The investigation of the SOX9 gene, responsible for the DSD phenotype in humans, revealed no significant structural variations such as deletions or duplications, suggesting that other genetic or epigenetic factors might contribute to the observed DSD phenotype in Brutus. Previous studies have associated the virilization of XX dogs with the SOX9 gene copy number and increased copy number in the CNV region upstream of this gene (27); however, these associations were not detected in our study. This implies that the DSD phenotype in Brutus may involve alternative loci, mechanisms, and/or genetic interactions.

The genomic sequencing of our French Bulldog along with the other three control dogs, including his father, has provided deeper insights into the genetic underpinnings of this DSD phenotype. Our analysis identified 43 SNVs unique to Brutus. Although these variants are classified with a modifier impact - suggesting they may not directly affect protein function - they could represent non-coding or regulatory variations potentially influencing gene expression and contributing to the observed phenotype. Notably, 8 of these SNVs are clustered within an 800 bp region on the X chromosome, specifically upstream of the spermiR sequence. SpermiRs are microRNAs critical for spermatogenesis and exhibit high and testis-specific expression patterns conserved across several mammalian species (28). This localization suggests that, while each SNP individually may have a minimal effect, their cumulative influence could be significant, potentially affecting spermiR’s gene regulation and, consequently, impacting the expression of downstream genes involved in fertility. This would explain the histological results obtained showing the lack of mature spermatozoa, the presence of an incompletely developed germinative epithelium, and Sertoli cells with nuclear alterations. Other significant discoveries made by ROH analysis provided further insights into the genetic architecture of Brutus compared to healthy controls. Brutus showed a significantly higher number of ROH segments and total ROH length, indicating a higher degree of homozygosity. This high level of homozygosity suggests significant inbreeding, contributing to the accumulation of homozygous deleterious alleles, which may be responsible for the observed phenotype.

The high F_ROH value, considering the total length of the autosomal chromosomes, suggests inbreeding, which may have exposed recessive alleles that contribute to the DSD phenotype. Inbreeding, while useful for maintaining breed purity, can significantly impact genetic health by increasing homozygosity, which in turn can correct both advantageous and deleterious traits within the population. Genomic inbreeding coefficients, which measure the proportion of an individual’s DNA inherited from recent common ancestors, provide a direct measure of inbreeding and are more accurate than pedigree-based estimates. This study shows that Brutus has a higher genomic inbreeding coefficient than control dogs, indicating a recent inbreeding event, most likely by mating between siblings. Typically, the inbreeding coefficient for an individual born from a sibling mating is around 0.25. However, in our analysis, Brutus exhibited an F_ROH of 0.31, which is higher than expected. This higher value is likely because the common ancestors of Brutus’ parents were not fully allozygous, indicating a certain level of autozygosity present in the breed. This is supported by the finding that all three individuals analyzed have comparable inbreeding coefficient values. To contextualize this value, we calculated the average inbreeding level of the breed by averaging the F_ROH of three normal genomes. We then estimated the theoretical inbreeding coefficient for an individual born from a sibling mating taking into account as a baseline level of inbreeding within the population the average value of the three normal individuals. Interestingly, our calculations yielded a theoretical value of 0.28 (0.29 when using the father of Brutus’s F_ROH value), which closely matches the empirically observed value. These findings support the hypothesis that Brutus is the offspring of related parents, specifically two siblings.

The findings from our ToppGene analysis provide valuable insights into the genetic underpinnings of the phenotype observed in Brutus. By examining the genes present within the ROHs in Brutus, we identified several significant associations, culminating in a total of 81 genes linked to our phenotype of interest. The presence of these genes in a homozygous state, likely due to recent inbreeding, could be implicated in a complex mechanism involving multiple pathways, all associated with sex development and hermaphroditism. Inbreeding can lead to an increased homozygosity of deleterious alleles, which can unmask recessive traits and disrupt normal developmental processes. Particularly noteworthy among these genes are SRD5A2, RSPO1, and SOX9. These genes have been highlighted in prior Pubmed publications as being associated with similar phenotypes in other species (29–31). The identification of these genes in our canine model suggests potential parallels in the genetic mechanisms underlying sexual development and hermaphroditism across species. This cross-species relevance underscores the importance of further investigating these genes in dogs.

In summary, our study underscores the complexity of disorders of sexual development (DSD) in dogs, highlighting the combination of chromosomal anomalies, absence of critical sex-determining genes, and unique genetic variants as underlying factors. This complexity is further compounded by the dual nature of inbreeding in breed management: while it aids in preserving breed-specific traits, it simultaneously elevates the risk of phenotypic abnormalities such as DSD. Future studies should focus on characterizing the specific mutations and their phenotypic consequences in the canine model, providing deeper insights into the genetic regulation of sexual development and contributing to our understanding of similar disorders in other species. Additionally, understanding the genetic basis of these conditions could lead to improved diagnostic tools and potentially inform strategies for managing or preventing such phenotypes in dog populations. Furthermore, future research should explore the regulatory mechanisms and potential epigenetic modifications that could elucidate the pathways involved in canine DSD, and consider the balance between maintaining breed purity and genetic health. Such investigations could ultimately pave the way for more effective approaches to breeding management and disease prevention in dogs.

### Supporting Information captions

**S1 Fig.** FISH analysis using an X-specific probe (CH82-201N14) in three healthy control dogs: (A) Brutus’s father, (B) Bufalo, and (B) Tauro. All three dogs exhibit a normal karyotype (78, XY), as indicated by the probe signal appearing on only one chromosome. Arrows point to the Y chromosomes in each panel.

**S2** **Fig.** 1% agarose gel electrophoresis showing the results of PCR performed with canine-specific SRY primers. Lanes are loaded as follows: Brutus’s father (A), Brutus (B), 2-log DNA marker (M; band sizes indicated in bp), Bufalo (C), and Tauro (D). The presence of the band in Brutus’s father, Bufalo, and Tauro, at the expected size (271 bp), indicates the presence of the SRY gene, while its absence in Brutus confirms the SRY-negative status.

**S1 Table:** Results of variant calling (performed with GATK) and subsequent annotation (executed with SNPEff). This table includes only those variants with call quality exceeding the filtering parameters described in the Materials and Methods section, which are not present in dbSNP and are unique to Brutus, with none of these variants found in heterozygous or homozygous form in the other three sequenced Bulldogs.

**S2 Table:** Detailed results of the Runs of Homozygosity (ROH) analysis for all sequenced samples (Tauro, Bufalo, Brutus, and Brutus’s father). Each column presents the following information: SAMPLE (identifier for each dog); CHR (chromosome number); POS1 and POS2 (start and end positions in CanFam6 of each ROH segment); KB (length of the ROH segment in kilobases); NSNP (number of SNPs present in the segment); DENSITY (density of SNPs in the segment) and PHOM (percentage of homozygosity within the segment).

**S3 Table:** List of genes annotated in CanFam4 within ROH fragments of Brutus and analyzed in ToppGene

**S4 Table:** The “Biological Process” results obtained from ToppGene

**S5 Table:** The “Mouse Phenotype” results obtained from ToppGene

**S6 Table:** The “Disease” results obtained from ToppGene

**S7 Table:** The “Pubmed” results obtained from ToppGene

**S8 Table:** List of genes found in ROH fegions of Brutus associated with sex development and hermaphroditism identified via ToppGene analysis

**S1 Text:** Clinical supplementary information

## Acknowledgments

We acknowledge financial support under the National Recovery and Resilience Plan (NRRP), Mission 4, Component 2, Investment 1.1, Call for tender No. 104 published on 2.2.2022 by the Italian Ministry of University and Research (MUR), funded by the European Union – NextGenerationEU– Project Title Telomere-to-telomere sequencing: the new era of Centromere and neocentromere eVolution (CenVolution) – CUP H53D23003260006 - Grant Assignment Decree No. 1015 adopted on 07/07/2023 by the Italian Ministry of Ministry of University and Research (MUR) (M.V.)

